# A revisit to a low-cost method for the isolation of microsatellite markers: the case of the endangered Malayan tapir (*Tapirus indicus*)

**DOI:** 10.1101/384651

**Authors:** Qi Luan Lim, Nurul Adilah Ismail, Ramitha Arumugam, Wei Lun Ng, Christina Seok Yien Yong, Ahmad Ismail, Jeffrine J. Rovie-Ryan, Norsyamimi Rosli, Geetha Annavi

**Affiliations:** Department of Biology, Faculty of Science, Universiti Putra Malaysia, Serdang, Selangor, Malaysia; China-ASEAN College of Marine Sciences, Xiamen University Malaysia; National Wildlife Forensic Laboratory (NWFL), Ex-Situ Conservation Division Department of Wildlife and National Parks (DWNP), Kuala Lumpur, Malaysia; Institute of Tropical Biodiversity and Sustainable Development, Universiti Malaysia Terengganu, Kuala Terengganu, Terengganu, Malaysia

## Abstract

There are many approaches to develop microsatellite markers. We revisited an easy and rapid Polymerase Chain Reaction (PCR)-cloning-sequencing method to design microsatellite markers for *Tapirus indicus*. Using six random amplified microsatellite (RAM) markers, this study had rapidly generated 45 unique genomic sequences containing microsatellites. After screening 15 terminal and seven intermediate microsatellite loci, we shortlisted five and seven which were amplified either by single- or multiplex PCR using the economical three-primer PCR method. Genotyping attempts were made with ten *Tapirus indicus* individuals using three of the terminal microsatellite loci and all seven intermediate loci. However, none of the terminal microsatellite loci were considered useful for population genotyping studies, while the seven intermediate loci showed good amplification but were monomorphic in the ten samples. Despite successful detection of amplified loci, we would like to highlight that, researchers who are interested in this alternative method for isolation of microsatellite loci to be cautious and be aware of the limitations and downfalls reported herein that could render these loci unsuitable for population genotyping.

## Introduction

Microsatellites, also known as short tandem repeats (STRs) or short sequence repeats (SSRs), are stretches of DNA consisting of tandemly repeated 1-6 nucleotides occurring at high frequency in the nuclear genomes of most organisms, with the length of a microsatellite locus typically in 5-40 repeats [1]. Slippage event during DNA replication is generally considered as the main mechanism for the expansion and contraction of microsatellites [2]. Mutation rates of the slippage events elevate from background rates when the repeat numbers in microsatellites are large [3]. Because of the hypermutability, microsatellite markers have been shown to be highly polymorphic [4]. They are widely used for molecular genetic studies including fingerprinting, parental or kinship analysis, population genetic structure and biological resources conservation [2,5,6].

There are various methods to develop microsatellite markers [7,8]. These methods have been improved over decades and are well described. An extensive review of microsatellite isolation methods has been published [7]. Some of these methods are still sufficient and successfully used in recent microsatellite marker development projects [9,10].

One of the conventional microsatellite isolation methods employs conversion of a random amplified microsatellite (RAM) marker, which is a multi-locus marker system, to one or more single-locus microsatellite marker. RAM was first described by Zietkiewicz et al. [11] and later further improved by Fisher et al. [12] to anchor the RAM primers consistently at the 5’ ends of two adjacent microsatellites. PCR amplification then produces amplicons that contain microsatellites at both ends of sequence. Extra microsatellite may also be present between the two termini of microsatellites. Next, specific primers can be designed for both the terminal and intermediate microsatellites. This technique has been used in the rapid generation of microsatellite markers in earlier studies [13–15].

*T. indicus*, or commonly known as Malayan Tapir or Asian Tapir, is an odd-toed ungulate (Order Perissodactyla) that occurs solely in Southeast Asia and also the only Old World extant member of Tapiridae family [16]. Its current population distribution includes regions of Myanmar, Thailand, Peninsular Malaysia, and Sumatra with not more than 2500 mature individuals worldwide, and it has been listed as ‘Endangered’ by the International Union for Conservation of Nature (IUCN) Red List [17,18]. The population genetic structure of this species in Peninsular Malaysia remains poorly understood, and this may potentially hamper conservation efforts (e.g. breeding management) in the nation. Genetic marker such as microsatellite has been used to aid conservation management in other tapir species [19], but microsatellites have not been developed or published for this spectacular species yet.

We followed the same technique as described in [20] with minor modifications and genotyped a few tapir individuals using microsatellite markers isolated from RAM marker. To our knowledge, our work was the first study to analyse microsatellites isolated with RAM markers on capillary electrophoresis system. We reported the potential of combining RAM or specific anchor markers and other techniques namely multiplex PCR and three-primer-method for genotyping. We also discussed the advantages and downfalls when using RAM markers in isolating microsatellites. Even though we did not detect any polymorphism in the genotyped samples through this protocol, we hope the reporting of the process and limitations that we had encountered would help other researchers with similar interest on this method to be cautious in the generation and interpretation of the data obtained.

## Materials and Methods

### Sample collection and DNA extraction

Whole blood samples were collected for microsatellite isolation from three *T. indicus* individuals kept in enclosures at the National Zoo of Malaysia or Zoo Negara (3°12’N, 101°45’E) and the Sungai Dusun Wildlife Reserve (3°40’N, 101°21’E) by the respective veterinary officers at the study sites and transported on ice to the laboratory at Universiti Putra Malaysia. All the sampling procedures were approved by the Institutional Animal Care and Use Committee, Universiti Putra Malaysia (ethical approval ref.: UPM/IACUC/AUP-R033/2016). For genotyping, blood samples on FTA^®^ cards (Whatman, UK) and tissue samples of ten putatively unrelated individuals were provided by the Department of Wildlife and National Park (PERHILITAN), Malaysia. Genomic DNA (gDNA) was extracted from these samples using the QIAamp^®^ DNA Mini Kit (Qiagen, Germany) following the manufacturer’s spin protocol. The gDNA was quantified by Quantus™ Fluorometer using ONE dsDNA dye (Promega, USA) and ranged from 0.2 to 100 ng/μL.

### Development of microsatellite markers

Six 5’-anchoring RAM primers selected from published literature (Table 1) were used for amplification. These RAM primers were selected based on the length of 5’-anchors (minimum 7 nucleotides). Single-primer PCR reaction mixtures contained final concentrations of approximately 10 ng of template DNA, 1× MyTaq Red Mix (Bioline, Germany), and 1.0 μM RAM primer, in a total volume of 25 μL. Two types of PCR reaction profiles were used: general and touchdown (PCR condition I in Table 2). Details on the PCR reaction profile and *Ta* used for each RAM primer are listed in Table 1. PCR amplification products were visualised on 2% agarose gel containing RedSafe™ Nucleic Acid Staining Solution. The remaining PCR products were purified using the Wizard^®^ PCR Clean-Up System (Promega, USA) or the QIAquick^®^ Gel Extraction Kit (Qiagen, Germany), following manufacturer’s instructions. The purified DNA amplicons were ligated and cloned.

**Table 1.**
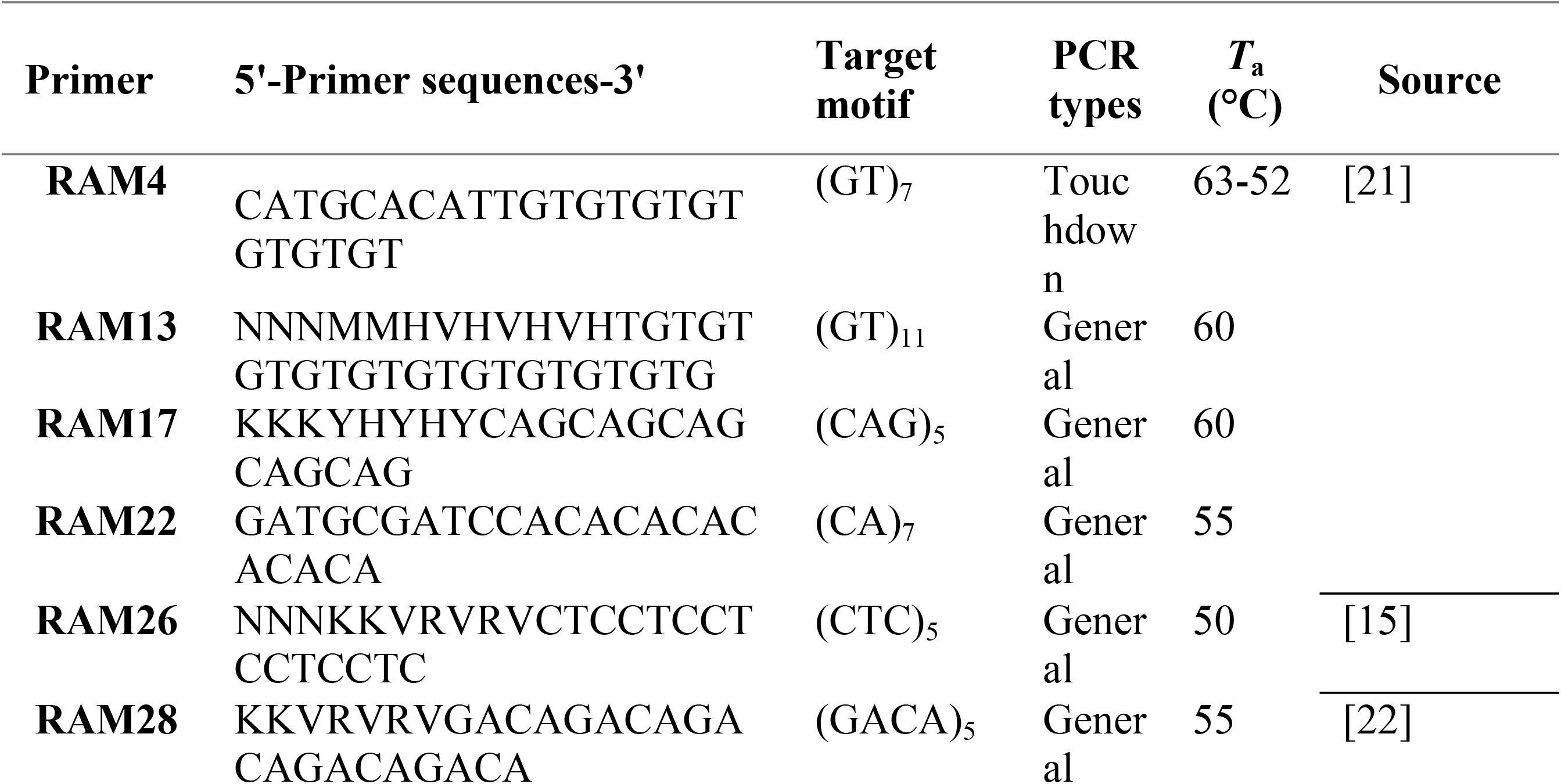
List of six primers tested for microsatellite markers isolation. Names given to the primers were used as a reference throughout the text.

**Table 2.**
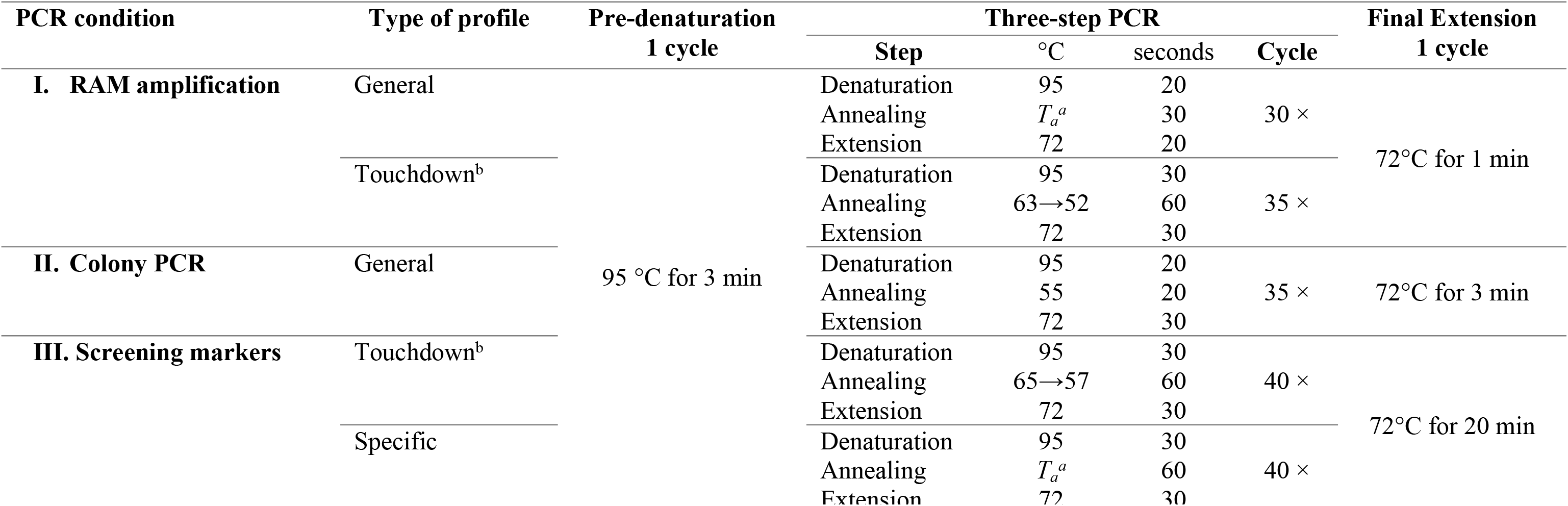
All PCR condition used in the development of microsatellite markers isolated from Random amplified microsatellite (RAM).

The purified PCR amplicons were individually ligated onto the pGEM^®^-T Easy Vector (Promega, USA) before transforming into JM109 High-Efficiency Competent Cells (*Escherichia coli*). Ligation reactions (in 10 μL volume per tube) for the products of each RAM were prepared with the following modifications to the protocol: 5 μL 2× Rapid Ligation Buffer, 0.5 μL pGEM^®^-T Easy Vector, 4.0 μL purified PCR product, 0.5 μL T4 DNA Ligase (3 Weiss units/μL). Then, 5 μL of ligation reaction was added to transform 20 μL of cells per tube, followed by 980 μL of super optimal broth (SOC) medium. The bacteria were cultured in an incubator at 37°C for 16-18 hours, plated on LB agar containing ampicillin (50 μg/mL) and X-gal (40 mg/mL), and placed at 4°C overnight before blue-white screening and colony PCR screening for insert sizes. Eleven to 35 white colonies on the agar plates per ligation reaction were randomly selected and subjected to colony PCR using the M13 primers (F: 5'-GTAAAACGACGGCCAGT-3'; R: 5'-AACAGCTATGACCATG-3').

Colony PCR was performed in reaction mixtures containing 1× MyTaq™ Red Mix, 0.25 μM of each primer in a final volume of 10 μL with PCR condition II (Table 2). The PCR amplicons were visualised on 2% agarose gel, and clones with different insert sizes were chosen for subsequent Sanger sequencing service on an ABI Genetic Analyzer ABI3730XL (Applied Biosystems, USA).

Sequences generated from the same locus were referred to as redundant sequences, and they were defined as sequences with ≥90% similarity and were detected by an online tool called ‘Decrease Redundancy’ [23] under default settings. From the output, non-redundant sequences generated from same RAM markers were paired sequence-by-sequence and compared using LALIGN [24,25] to detect highly similar sequences or chimeras. Sequences showing possible chimeric structure between two different sequences were discarded.

Perfect microsatellites present at both ends (terminal microsatellites) and within the sequences (intermediate microsatellites) were detected using IMEx [26] using the following criteria: minimum eight repeats for mononucleotide , four repeats for di- and trinucleotide, three repeats for tetranucleotide, and at least two repeats for penta- and hexanucleotide repeat motifs. Compound microsatellites constituting two or more repeat motifs within 10 bp distance apart were also detected. Interrupted microsatellites were detected by manually examining the upstream and downstream segments of individual perfect microsatellites for any repeats of the same motif within 10bp distance.

Unlike usual primer pair design, primer pairs of the isolated loci carrying terminal microsatellites constituted a non-degenerate RAM or specific anchor primer pairing with a specific primer (termed *internal primer*) designed from intermediate sequence in between two terminal microsatellites. A specific anchor primer has the sequence of the targeted repeating motif and a specific 5’ anchor that corresponds to its RAM marker. General criteria of primer pair design using Primer3Plus ([27]). The designed primers are listed in S1 Table.

#### Screening of markers

To test the designed primer pairs, PCR amplification was performed in 10 μL single-plex PCR reaction mixtures containing 1× MyTaq Red Mix (Bioline, Germany), 2-20ng of template DNA, and 0.4 μM of primer mix using touchdown profile of PCR condition III (Table 2). Primer pairs of loci that produced bands of expected sizes with minimal unspecific products were chosen for further analysis by fragment analysis (Table 3), and their internal primers were redesigned to attach at the 5’ end one of the following sequences: M13 (5’-6-FAM-TCCCAGTCACGACGT-3’), Neomycin rev (*5’-6-FAM-* AGGTGAGATGACAGGAGATC-3’), M13modB (5’-tfEX-CACTGCTTAGAGCGATGC-3’), Hill (5’-tfEX-TGACCGGCAGCAAAATTG-3’) or T7term (5’-ROX-CTAGTTATTGCTCAGCGGT-3’) for subsequent three-primer PCR reactions [28–30]. The tailed internal primer had primer length up to 46 bp.

**Table 3.**
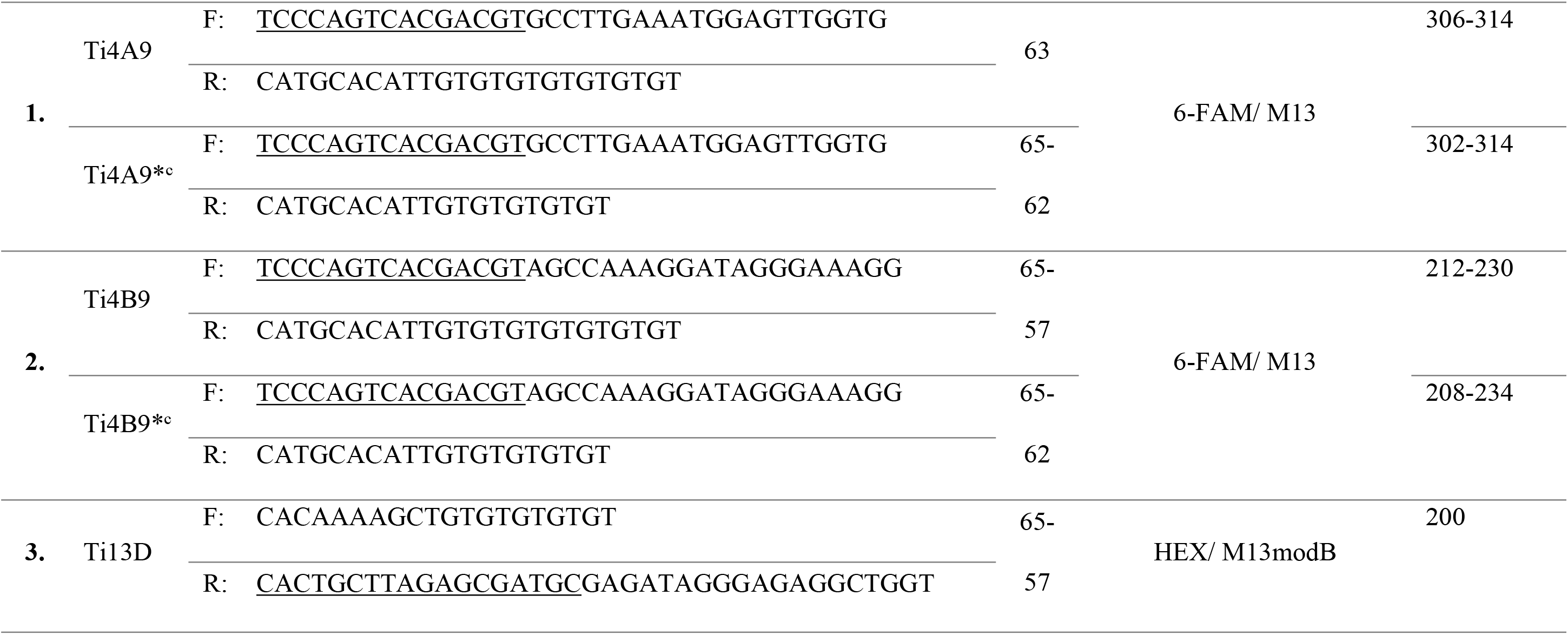
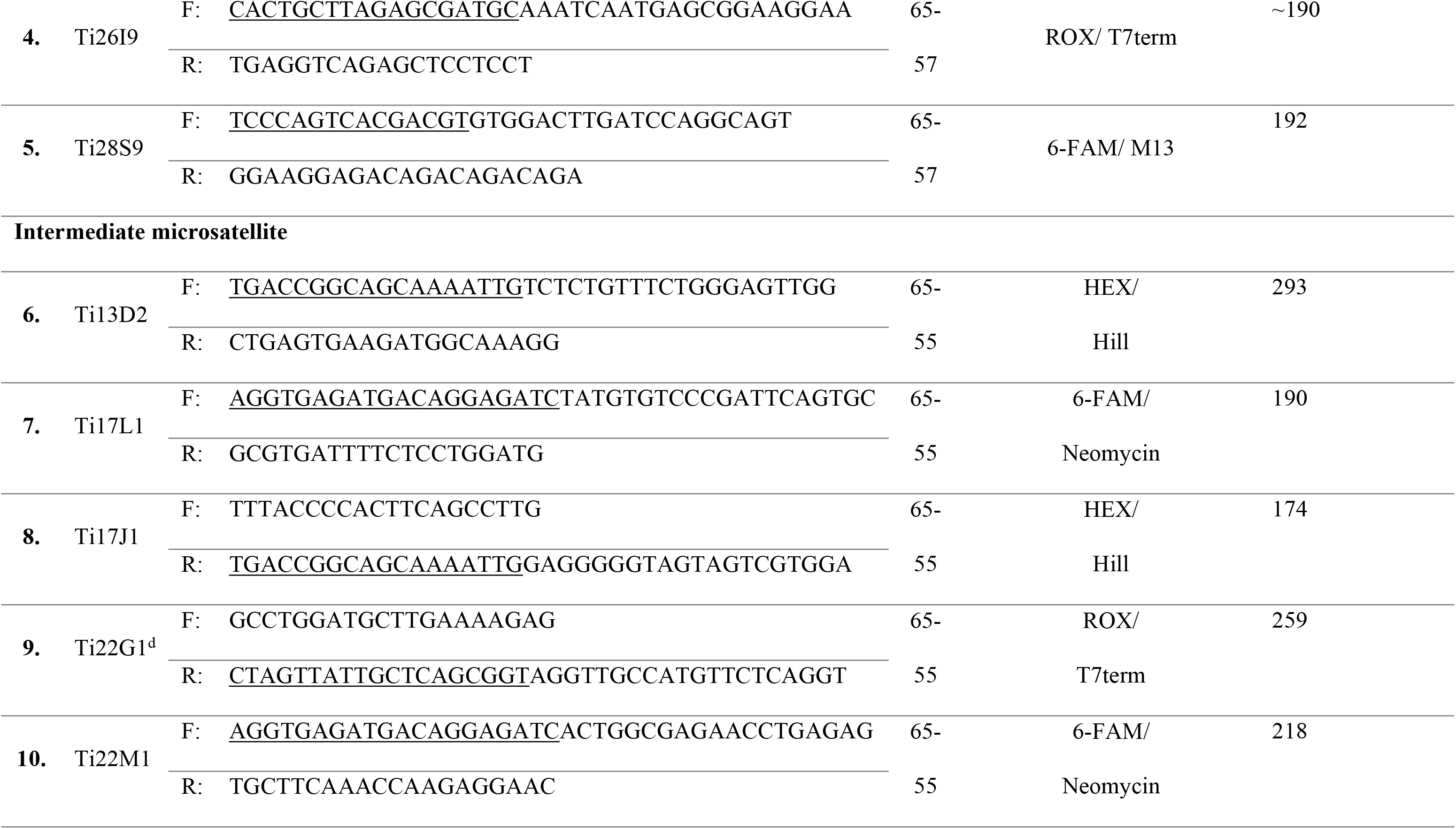
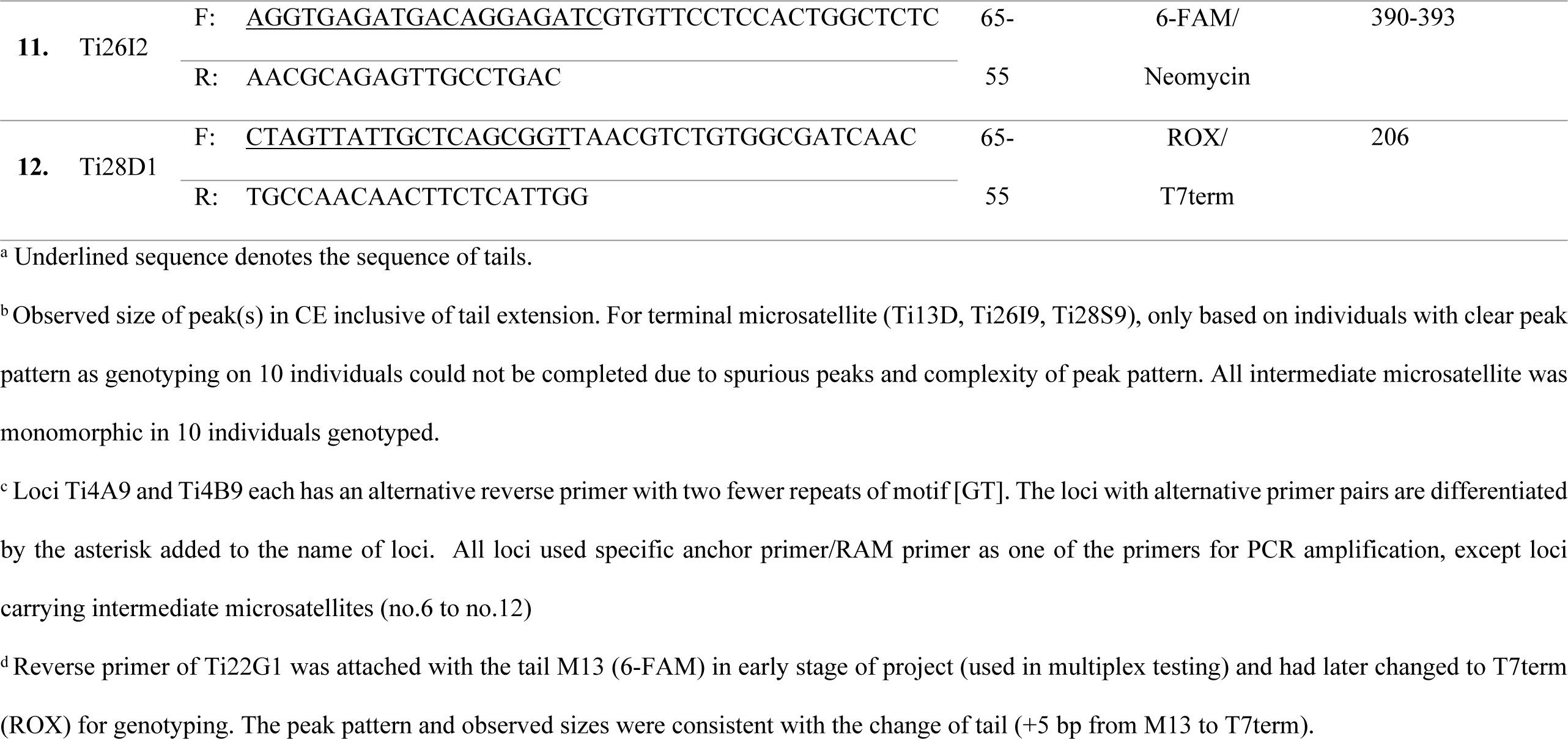
List of 14 primer pairs from 12 loci developed from RAM markers and the annealing temperature (*T_a_*), observed and expected sizes. The primers were extended with tails that were customised for a dye colour. CE – capillary electrophoresis.

Primer mix containing three primers in the ratio of 4:1:4 (final concentration of 10 μM: 2.5 μM: 10 μM) for nontailed internal primer/ specific anchor primer/ RAM primer: tailed internal primer: and fluorescent-labelled tail respectively, were prepared. PCR amplification was run in 20 μL single-plex PCR reaction mixtures containing 1× MyTaq Red Mix, 0.4-1.0 μL primer mix, and 2-20 ng of template DNA. PCR was run using PCR condition III (Table 2) at specific *Ta* listed in Table 3.

Concurrently, another PCR amplification was performed that excluded the fluorescent-labelled tails and contained only a non-tailed internal primer/ specific anchor primer and a tailed internal primer in equal concentrations of 10 μM, to test the possible inhibitory effect of fluorescent-labelled tail primers.

To test the compatibility of the designed primers in multiplex PCR, Ti4A9 and Ti4B9 (Table 3) were multiplexed together using touchdown profile of PCR condition III. Another multiplex PCR including loci Ti13D, Ti22G1 and Ti28S9 were run in the same conditions, except that 0.4 μL, 0.2 μL, and 0.3 μL primer mix of the loci in same order were added. The PCR products were sent for fragment analysis on ABI3730XL.

We also included all seven intermediate microsatellites markers and only three terminal microsatellite loci (Ti13D, Ti26I9, and Ti28S9) for genotyping 10 individuals of *T. indicus* in single-plex PCR with 1.5-3 ng DNA template using the same PCR condition III and sent for fragment analysis. Genotypes were examined and scored using R package *Fragman* in R Studio 1.1.442 [31–33]

## Results and Discussion

### Isolation of microsatellite loci

This study attempted to isolate microsatellite loci from a mammalian genome with RAM primers. Previous studies have succeeded with the use of this technique in plants, fungi, and insects [15,21,34,35]. The use of RAM amplification, which obviates the steps of genome fragmentation and enrichment. Instead, the simple workflow allowed rapid generation of microsatellite-containing genomic sequences with an affordable budget, less work, and in a short time (<2 weeks per isolation).

The recovery of recombinant inserts from 135 screened colonies was 98.5% (1.5% were false positives). Sixty-five positive clones were sent for sequencing, and all the sequenced fragments carried at least two microsatellites. Among these sequences, 18 (27.7%) were regarded as redundant sequences based on 90% max similarity search. One unique sequence was discarded for possible chimaeric structure. Another sequence was discarded due to short length (187 bp) and hence for primer design.

The remaining 45 sequences were deposited in GenBank (accession numbers.: MH292918 - MH292962). From these sequences, a total of 119 non-interrupted, perfect microsatellites were detected by IMEx [26] under our selection criteria (see Methods), and more than half of them (n = 68) were from the termini of each sequence. In addition, 20 compound microsatellites and six interrupted perfect microsatellites were detected. The intermediate microsatellites obtained were typically low in repeat number: six to 11 repeats for mononucleotide motifs, and at most four repeats for dinucleotide motifs, six repeats for trinucleotide motifs, three repeats for tetranucleotide motifs, three repeats for pentanucleotide motifs, and two repeats for hexanucleotide motifs.

More repeat number found in the terminal microsatellite loci than in their respective RAM primers may be an indicator for correct hybridisation of the 5’-anchor to its complementary template (Fisher et al. 1996). In our study, 69.7% of 89 detected terminal microsatellites have repeat number equal to their respective RAM primers. Comparing to three other studies that reported equal repeat number using similar method, the proportions of loci with equal repeat number to RAMs were 35% (60 loci; [13]), 50% (26 loci; [12]), and 60% (10 loci; [35]). The number in parenthesis is the total number of terminal microsatellites that were isolated regardless of how many different 5’ anchored RAM used. We recorded the highest percentage of such loci largely because three of the RAMs (RAM17, RAM26, and RAM28) all yielded tri- or tetranucleotide microsatellites with five repeats. For the other three RAM markers, half of the terminal microsatellites yielded had the same repeat number in the RAM primers.

### Testing of designed primers

In this study, 24 primers pairs were designed (S1 Table). To test the terminal microsatellites, we selected 15 loci comprising of: four loci from RAM4, eight loci from RAM13, one locus each from RAM17, RAM26 and RAM28. Two loci from RAM4 had their primers redesigned to test under different conditions, adding to a total of 17 primer pairs tested for terminal microsatellites. To test the intermediate microsatellites, we selected seven loci comprising of: two loci from RAM17, two loci from RAM22, one locus each of which from RAM13, RAM26 and RAM28.

Of the 24 tested primer pairs, Ti4A9, Ti26I9 and all intermediate microsatellite loci each yielded a clear single band of expected size, while the other eight primer pairs yielded expected bands with some degree of unspecific amplification. The rest either failed in amplification or yielded multiple unspecific bands (descriptions in S1 Table). Expected product sizes of all 14 primer pairs in Table 3 were detected in capillary electrophoresis.

Generally, loci amplified by specific anchor primers produced less stutter peaks than using RAM primers based on the observations on TI4A9-Ti4A9* and Ti4B9-Ti4B9* pairs (Fig 1). When the motif repeat in RAM4 primer was reduced by two repeats, it was expected that the stutter bands (due to primer slipping to 3’ end) could be reduced by increasing the proportion of specific anchor (10 in 24 bases to 10 in 20 bases) to improve the priming efficacy of the 5’-anchor. Both loci, Ti4A9* and Ti4B9*, have shown such improvement but in the expense of yield and increased risk of unspecific amplification that led to noise (Fig 1B, 1C).

**Fig 1.**
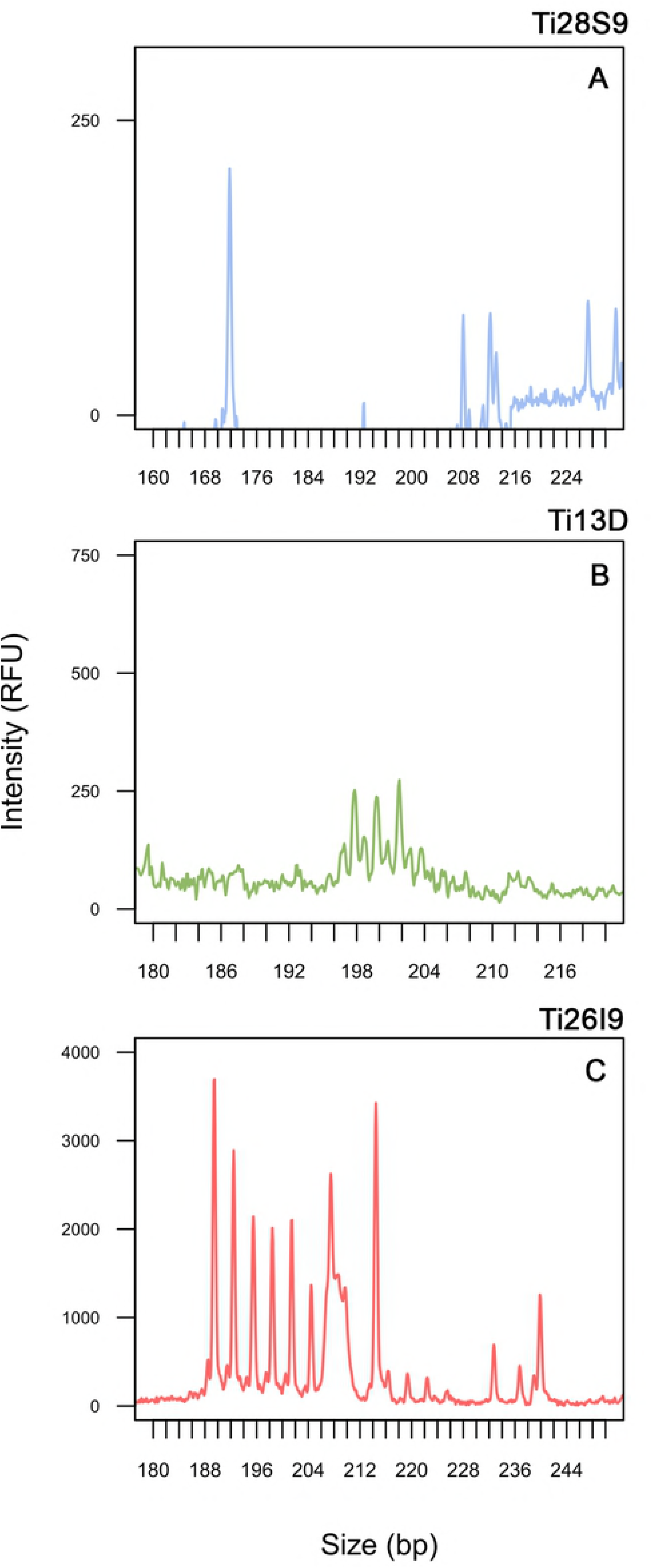
Genotype for the tested five loci from Peak Scanner 2.0. (A) Multiplex PCR products of Ti4A9 and Ti4B9 at final dilution factor (FDF) of 100. (B) Co-loading at 60 FDF of two single-plex PCR products, the Ti4A9 (63^°^C, 40 cycles) and the Ti4B9* (65-62^°^C, 40 cycles). (C) Co-loading at 60 FDF of two single-plex PCR products, the Ti4A9* (65-62^°^C, 40 cycles) and Ti4B9* (65-62^°^C, 40 cycles). (D) and (E) Multiplex PCR products of 3 loci namely Ti22G1, Ti28S9, and Ti13D. Some undesired signal in the result were: **Ar –** Dye artefact. Template DNA was extracted from blood of a single tapir individual except for Ti4B9* in (C), and 3 loci in (D) and (E). Ti4A9* and Ti4B9* used shortened RAM4 (20 bases) as the reverse primer, while Ti4A9 and Ti4B9 used original RAM4 primer (24 bases) as the reverse primer. Unless otherwise specified, PCR for all shown result was touchdown profile of 65-57°C for 40 cycles.

### Inhibitory effect of fluorescent-labelled tail

During optimisation, one of the loci, Ti26I9 failed to be amplified when M13modB was used as the labelling tail primer but worked when the tail was removed from the PCR reaction, which was repeated in three PCR trials. This observation suggested an inhibitory effect of fluorescent-labelled tail primer used in the three-primer method. The primer pair (originally F: 5’-CACTGCTTAGAGCGATGCAAATCAATGAGCGGAAGGAA-3’ and R: 5’-TGAGGTCAGAGCTCCTCCT-3’) was then redesigned to the one listed in Table 3. All the other 13 primer pairs in Table 3 were successful in amplifying their expected products by both the three-primer PCR and normal PCR (excluding tail primers and using equal concentration of forward and reverse primers).

### Multiplex PCR and genotyping

Multiplex PCR involving two loci (Ti4A9 and Ti4B9), that shared a common RAM primer, a tail sequence attached to their internal primers and a fluorescent-labelled tail, was feasible. Both loci were amplified and detected during fragment analysis (Fig 1A). Another multiplex PCR involving two terminal microsatellite loci (Ti13D and Ti28S9) and one intermediate microsatellite locus (Ti22G1) was also successful. The compatibility may be attributed to the different repeat motifs carried on the primers i.e. (TG)_5_ on Ti13D and (CAGA)_3_ on Ti28S9, while Ti22G1 comprised of a pair of specific internal primers. Multiplexing primers carrying same or complementary repeat motifs may result in increased formation of primer dimer leading to low efficiency or inhibition of PCR.

Genotypes of ten individuals by all seven intermediate microsatellites markers were overall monomorphic, except for locus Ti26I2 (Table 3). However, due to a spurious peak at 3 bp distance away from the main peak we could not confidently score the genotypes even though consistent peak patterns could be obtained from different runs of PCR using the same DNA samples. Fig 2 shows an example electropherogram for the three terminal microsatellite loci used for genotyping. We were also not able to complete the genotyping of 10 individuals using the three terminal microsatellite loci due to dubious specificity (Ti28S9, with an extra, consistent peak near 400bp), complicated peak patterns accompanied with noise (Ti13D and Ti26I9), rendering them not suitable for further analysis.

**Fig 2.**
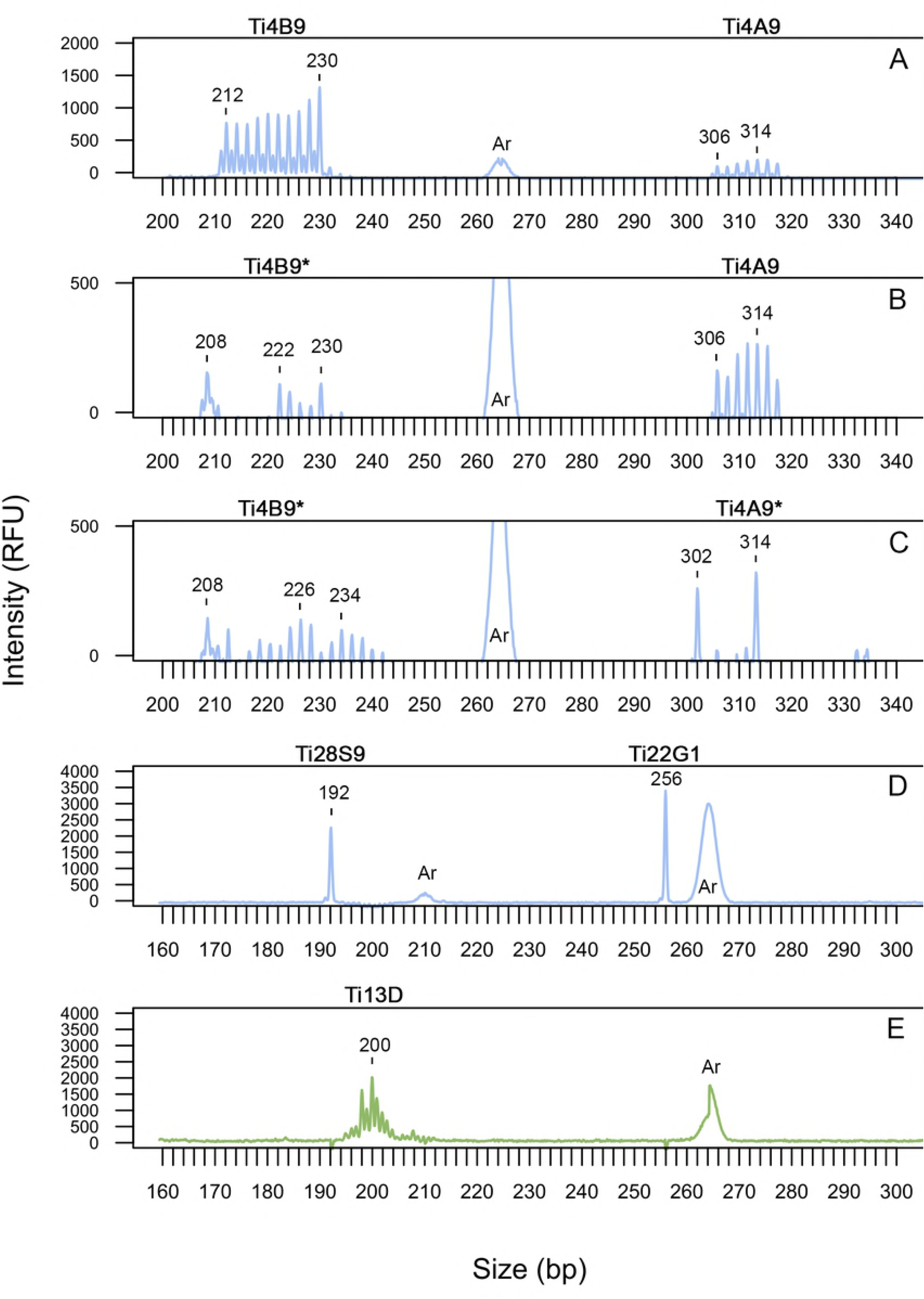
An example of electropherogram of three terminal microsatellites fragments amplified. (A) Ti28S9, (B) Ti13D and (C) Ti26I9. Final dilution factor (FDF) in the range of 5-12. All was amplified in single-plex PCR by touchdown profile (65-57°C, 40 cycles)

Although specific anchor primers of terminal microsatellite markers had shown successful amplification of targeted loci and applicable for genotyping on gels [13–15], none of these studies genotype the samples by capillary electrophoresis. From the result, we have shown that even though microsatellite markers isolated using the protocols described could be detected in the fluorescent capillary electrophoresis, the quality of peak pattern varied across samples. On the contrary, amplification of intermediate microsatellite loci using specific internal primers did not encounter such issue.

### Limitations

The nature of primer design for terminal microsatellites (that must include a length of repeat motifs and a 5’ flank derived from RAM primer) has rendered them to be prone to random amplification of unspecific products. The 5’ anchors of RAMs used in this study to develop microsatellite markers were 7-12 bases including 5-9 blocking bases to prevent slipping of primer to 3’ end [12]. However, the sequences of anchors used for primer design were not always in perfect match to their template, depending on the stringency of RAM amplification. Too high a *Ta* may produce low yield, while low *T_a_* may allow mismatches at the 5’-anchor. This was evident by the nucleotide mismatches seen in the 5’-anchors between pairs of sequences that were supposedly amplified from the same loci during RAM amplification. The mismatch ranged from 0-9 nucleotides with an average of 4 nucleotides (or an average of 57% similarity between 5’-anchors of the pairs of sequences). In extreme cases only 1-2 nucleotides in the anchors were shared.

Even though it was suggested that reproducibility could still be achieved in the presence of primer-template mismatches [36], the amplification specificity may be further compromised by the presence of microsatellite repeats at the more critical 3’ end of primer. In fact, the intermediate microsatellites produced a cleaner electropherogram with minimal noise peaks and higher relative fluorescence units (RFU) compared to terminal microsatellites. Furthermore, there was only very limited number of nucleotide usable for primer design. As observed in some of the flanks (found in <10% of genomic sequences), truncation of one or more nucleotides from the 5’-ends of anchors, had made primer design even less flexible.

Other than null allele [13], mis-scoring of fragment size may arise from hybridisation between RAM primer and template carrying less repeats than the one present in RAM. For instance, the smallest peak of Ti4A9* (302bp) was four bases less than the one detected in Ti4A9 (306bp), which corresponded to four-base reduction in the modified RAM4 primer ([GT]_7_ to [GT]_5_).

In this study, we managed to develop six amplifiable microsatellite markers for *T. indicus*, although they were monomorphic in ten individuals, which may be due to low genetic diversity in this species. Genotyping a larger sample size in the future may reveal useful and polymorphic markers. In general, we do not recommend using RAM markers for microsatellite isolation due to a few limitations discussed above.

## Acknowledgements

We would like to thank Dr. Donny Yawah (PERHILITAN), Dr. Mat Naim Bin Haji Ramli (National Zoo of Malaysia) and Dr. Kavitha Jayaseelan (National Zoo of Malaysia) for their assistance in collecting blood samples at the Sungai Dusun Wildlife Reserve and the National Zoo of Malaysia respectively. We also thank Mdm Noor Azleen binti Mohd Kulaimi (PERHILITAN) for aiding in the collection of tapir samples from the National Wildlife Forensic Laboratory, PERHILITAN.

## Supporting Information

**S1 Table. List of all designed primer pairs of 15 terminal and 7 intermediate microsatellite loci**. The primers were isolated using six random amplified microsatellite (RAM) markers. Expected size and description on the specific anchor primer (SAP) and band pattern on gel is provided. CE - capillary electrophoresis.

